# QUANTITATIVE ETHNOMEDICINAL SURVEY OF SHRUBBY PLANTS, USED BY THE LOCAL COMMUNITIES OF DISTRICT LAHORE (PUNJAB), PAKISTAN

**DOI:** 10.1101/2020.06.18.158865

**Authors:** Shabnum Shaheen, Sobia Sarwar, Nidaa Harun, Sana Khalid, Khadim Hussain, Farah Khan

## Abstract

**Background:** Indigenous knowledge of medicinal flora is a baseline for the production of plant based commercial drugs. Current study was planned to explore the ethnomedicinal uses of shrubs in traditional health-care system of District Lahore. This study also aimed to represent the conservation status of these natural resources which are decreasing day by day due to their overexploitation and deforestation.

**Methods:** The key informants were identified by employing the snowball technique. Data was collected by conducting semi-structured interviews with 103 informants from different localities (herbal markets, nurseries, gardens) of district Lahore. Collected data about medicinal shrubs were analysed on different data analyses parameters such as relative frequency of citation (RFC), use value (UV), informant consensus factor (ICF), fidelity level (FL) and relative importance (RI). In addition, SPSS 22 software was used for statistics analysis and interpretation of associations among different studied parameters.

**Results:** A total of 115 ethnomedicinal shrubs belonging to 50 families were reported to be used against different diseases. The study area was dominated by Fabaceae family (n=10). The RFC ranged from 0.02 (*Deutzia scabra* and *Euonymus japonicus*) to 0.85 (*Rosa indica*) while UV ranged from 0.01 (*Cestrum diurnum* and *Garcinia aristata*) to 0.23 (*Jasminum grandiflorum* and *Hamelia patens*) and RI ranged from 0.16 (*Garcinia aristata*) to 43.73 (*Tabernaemontana divaricata*). Moreover *Lawsonia inermis, Piper nigrum, Punica granatum, Rosa indica* and *Vitis vinifera* reported with 100% FL whereas maximum ICF were calculated by gastrointestinal diseases i.e., 0.45. On the basis of cluster analysis ethnomedicinal shrubs were categorized into two groups i.e., high valued and low valued. It was also found that most of the species in high valued group (n=29) were rare in study area due to their over exploitation. This study also documented 104 new use reports for ethnomedicinal shrubs.

**Conclusions:** This study documented significant indigenous knowledge about ethnomedicinal shrubs used by the local people of District Lahore. This knowledge could be worthwhile in discovering and developing new plant-based drugs. Apart from this, current study also revealed that most valuable medicinal species are declining in their number due to over usage and mismanagement. Conservation strategies for medicinal plants of District Lahore are highly recommended.

## Background

By definition shrubby plants are regarded as those woody plants which are usually smaller than trees and consisting of numerous main stems arising at or nearby the ground level. These shrubs play significant ecological and commercial roles in our lives, such as they help to prevent soil erosion, support in beautifying our landscapes, used as a fuel wood and lumber resource and even many of them work as a spring of foodstuffs for humans as well as for diverse range of animals. Apart from these uses, these shrubs also hold a significant medicinal value and serve as major therapeutic agent in cure of multiple disorders such as Butea monosperma (Lain.) is used to treat menstrual pain, Helicterus isora (Linn.) reported to treat colic and intestinal disorder whereas Rosa canina (*L.*), Helichrysum stoechas (L.) and Myrtus communis *(L.)* used against obesity.

The ethnomedicinal value of shrubby species is well supported by various ethnobotanical studies around the world. In Iran 16 medicinally important shrubs were identified, which were renowned as antipyretic, anti-diarrheal, anti-inflammatory, laxative, blood purifiers and for cure of toothache. In another ethnobotanical survey conducted in Talfila region of Jordan, reported 33 shrubs species with medicinal potential. Similarly an ethnomedicinal survey in Eastern Cape Province of South Africa designated 25 shrubs in treatment of multiple urogenital diseases. Additionally in two different studies led in Ethiopia (Meinit ethnic group) and Nepal (Mid-Hills) stated 13 and 36 shrubs with medicinal efficacies respectively. Moreover in recent studies conducted in China (Maonan people) revealed 68 medicinal shrubs whereas in India: a total of 26 medicinally important shrub species belonging to 19 genera and 16 families were recorded from the region of Kashmir.

In Pakistan there are almost 400–600 medicinal plants due to its unique and diversified climatic conditions, ecological and topographical zones. A lot of ethnobotanical studies in various geographical regions of Pakistan had been conducted and these studies reported significant number of shrubs for their medicinal value. An ethnobotanical study in the region of Swat, recorded 106 plant species out of which 16% were shrubs which are used to treat many disorders such as gastro-intestinal disorders, dermatological and topical diseases, urinary complaints, respiratory illness and dental problems. In a similar study conducted in District Karak, Pakistan about 18 shrubby species used by indigenous people for the cure of various ailments [16]. In a FATA survey it was found that among the indigenous communities of Mohmand Agency, 9 shrubs were well famous in treatment of various infections. Furthermore in an ethnomedicinal study of the flora of Karakoram-Himalayan range, Pakistan, reported 13 various shrub species for management of common ailments. As traditional knowledge on herbal medicines usually transfers from one generation to another generation orally this may be a cause of information gap by passing time. So, there is a dire need to preserve the information in written form. Furthermore, the unregulated use of these wild resources could lead to the threatening of species in their particular habitats. Therefore the relative abundance status of these shrubs must be evaluated by consistent intervals. Hence, the current endeavour focussed to collect and document the ethnomedicinal data about the shrubs inhabited in district Lahore (Punjab) Pakistan. This will provide the baseline data for future phytochemical and pharmacological studies.

The main objectives of this survey were to document the (i) all ethnomedicinal shrubs of study area (ii) use part and the use pattern of the plants, diseases cured and (iii) other ethnobotanical uses of enlisted medicinal shrubs (iv) high and low valued ranking groups of enlisted ethnomedicinal shrubs and (v) the relative abundance value for evaluating their conservation status in study area.

## Methods

### Study area

Present study was carried out in premises of Lahore city that lying between 74° 20’ 37” E and 31° 32’ 59”N covering an area of 404 square kilometres. On the north and west of it is bounded by Sheikhupura District; on the east there is Wagah and on the south thers is Kasur District, whereas Ravi River flows at northern side of Lahore. Climate of this city is semi-arid, hottest months are June and July with temperatures ranging between 40°C-48°C, the coldest months are December and January where temperature range is from 4-6°C. Due to the diverse climate range it has wide variety of flora and fauna both natural and cultivated. For the data collection about shrubs different nurseries (Model Town plant nurseries, Abbas nursery, Faizan nursery form, Ishtiyaq nursery, Plant nursery and Al-Madina nursery), gardens (Jilani Park, Lawrence Garden, Jallo Park, Safari Park, Model Town Park) and Botanical garden of Lahore College for Women University, Govt. College University and Punjab University, Lahore) had been selected randomly from north, south, east and west coordinates of Lahore (Fig 1). However their medicinal efficacy was inquired from herbal practitioners by visiting major herbal markets (Akbari mandi and Asghari mandi) of Lahore.

**Fig.1.**
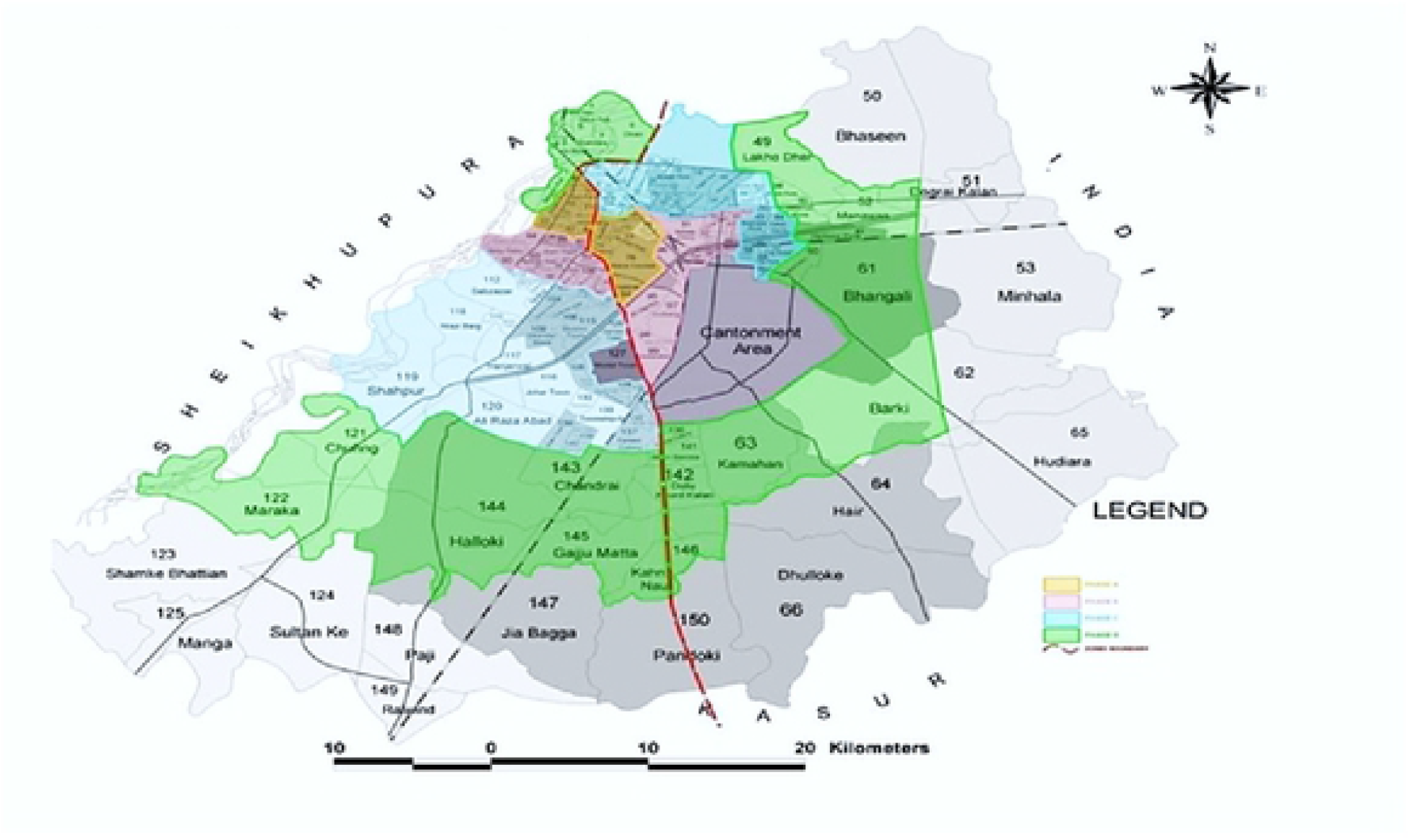
Map of study area

### Ethnomedicinal Inventory

Before collecting the data, code of ethics as set by ‘International Society of ethnobiology’ was strictly followed. Formal ethical permission was also taken from the chairperson of local government and local informants of study areas. The data were collected between the months of March 2017 to December 2017 from the different targeted areas of Lahore. For data collection, 103 respondents were interviewed and key informants were gardeners, nursery vendors, local people and herbalists (Table 1). Data about the shrub species was gathered by questionnaire and grouped discussions. Questionnaire and interview consisted of two parts; first part was about the data about the informant such as name, age, gender, education and profession. Second part included the information regarding to shrub medicinal use i.e. range of diseases that can cure by particular shrub, part used for medicinal purpose and any other uses apart from medicinal purpose. Most of the information was reported in Urdu and in vernacular language of area i.e. Punjabi. However, later on this data was translated back into English for data analysis ease and documentation purposes.

### Collection of shrubs and identification

Several site visits were made with some knowledgeable resource persons (gardeners or nursery care takers) for the identification and collection of shrub samples. The particulars of each shrub specimen i.e. collection date; habitat, local names, habit and flowering periods were also noted throughout each site visit.

After collection, plant species were pressed dried, sprayed with a preservative 1% HgCl_2_ solution and mounted properly on herbarium sheets. Preliminary identification of the collected shrubs was done by the expert taxonomist available in botany department of Lahore College for Women University (LCWU); Lahore, Pakistan. Whereas later on herbarium of Quaid-i-Azam University, Islamabad, Pakistan and National Agricultural Research Council were also consulted for further taxonomic identification. Naming of plant species and their family was confirmed by International Plant Name Index (IPNI) [25]. Voucher numbers were assigned to each specimen and deposited at the herbarium of the Department of Botany, LCWU Lahore, Pakistan.

### Estimation of relative abundance

For the determination of relative abundance, visual assessment was used. In this method, randomly sites were selected from study area and the presence of each species was counted and recorded, and then percentage relative abundance was calculated by using the following formula;

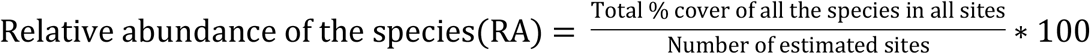

Species were grouped into different groups i.e. Abundant, Common, Frequent, Occasional and Rare (ACFOR) by using relevant scales of abundance (Table 2). Abundant indicates that species can be seen in large number. Common means species can be seen in appropriate season. Frequent means species were not seen consistently during visiting of a specific site. Occasional means species are present in restricted area and there number is less. Rare indicates even less probability of occurrence.

### Data analysis

All data was recorded by using Microsoft excel 2013. Ethnobotanical data was analysed by several quantitative parameters such as Relative Frequency of Citation (RFC), Use Value (UV), the Informant Consensus factor (ICF), Relative importance (RI) and the Fidelity Level (FL).

#### a) Relative Frequency of Citation (RFC)

The Relative Frequency of Citation (RFC) determines the relative importance of plant species based on a number of informants for each species and total informants in the study. It was calculated by dividing “FC” by the total number of informants in the whole survey (N) [28].

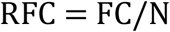

Where, FC stands for the frequency of citation. (Number of informants of a particular species)

#### b) Use value (UV)

The used value shows the recorded used reports of plants species by indigenous people. It tells about the relative importance of use of plant species based on the total number of informants. It was calculated by following formula:

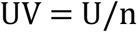

Where “U” refers the total number of uses per species while “n” is the number of informants who reported on the plant species.

#### c) Informant Consensus Factor (ICF)

Use of plant species to cure different disease types is determined by Informant consensus factor. It is calculated by the following formula.

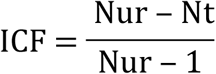

Where “Nur” is the number of use reports to treat a disease category, and “Nt” is the number of plant species used for treating that disease category.

#### d) Relative Importance (RI)

The relative importance (RI) describes the importance of use of each plant species and body organ systems treated by it. It is calculated as.

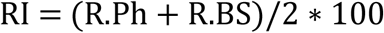

Where, “R. Ph” stands for relative pharmacological properties. “R.Ph” is calculated by dividing the number of uses (U) with the total number of use reports in the whole study. “R.BS” stands for relative body systems treated. “R.BS” is calculated by dividing the number of body system treated by a plant species with the total number of body systems in the whole study.

#### e) Fidelity Level (FL)

Fidelity level determines the percentage preference of a specific use of each plant species over other species. It is calculated by using formula.

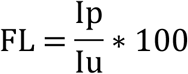

Where “Ip” is the number of informants who told about the use of a given species for the treatment of a specific disease and “Iu” is the total number of all informants who reported all uses about given plant species.

#### f) Statistical inference through Statistical Package for the Social Sciences(SPSS)

For making groups of high and low valued groups of ethnomedicinal shrub species, Hierarchical Cluster Analysis (Squared Euclidean distance method) in the SPSS 23 software was applied. Moreover descriptive statistical analysis (frequency and cross tabulation) was also employed to find out the association between different parameters of the survey.

#### g) Graphical illustrations

Microsoft Excel (2010) was used to convert selected data items into different types of graphical illustrations.

## Results and Discussions

### Demography of informants

Total 103 respondents took part in this study, out of which 77.67% were men and 22.23% women, whose ages ranged from 20 to 89 years. The smaller number of female informants showed the cultural and social limitations of the study area where females were found reluctant in expressing their views. Among the total informants: herbalists were 29.12%, gardeners 38.83% and 32.04% were local people (Table 1). In concern with education level it was observed that 47.57% informants were primary pass (5 years), 27.18 attended middle school (8 years), 14.56% secondary pass (12 years) and 10.68% had higher degrees. These educational facts declared that mostly the people of study area were educated and this is because of high literacy rate in District Lahore i.e., 64.7%.

### Diversity of ethnomedicinal shrubs

During ethnobotanical study of shrubs of Lahore, 115 shrub species belonging to 50 families from different localities were reported. Maximum number of species belonged to family Fabaceae (n=10) followed by Euphorbiaceae and Lamiaceae (n=9) and Apocynaceae (n=8). Greater number of Fabaceae members is probably due to the fact that Fabaceae is the third largest family in Pakistan. Moreover this family (Fabaceae) is also famous for its medicinal worth. Whereas Family Lamiaceae also reported for its antiseptic, anti-inflammatory and antimicrobial properties.

**Fig2.**
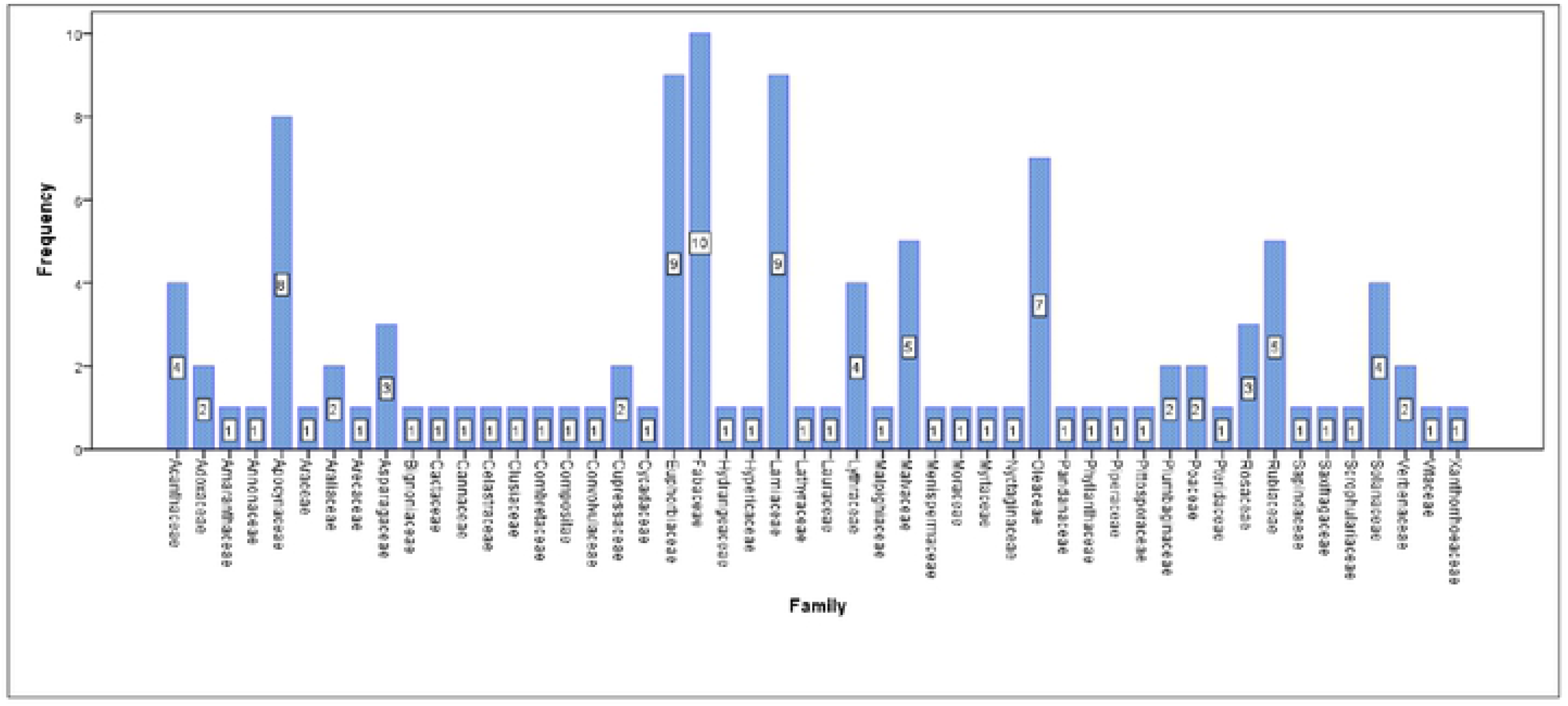
Numbers of ethnobotanical shrubs in each family.

### Used part of ethnomedicinal shrubs

From the descriptive statistics (frequency calculation) it was observed that nearly all parts of shrubs used for the medicinal purpose. Therefore local people reported whole plant usage is maximum i.e. 24.43%. Leaves (22.73%) were the second most highly used part followed by roots, flowers, seeds, fruits, bark, stem and plant sap (Figure 3).

**Fig 3:**
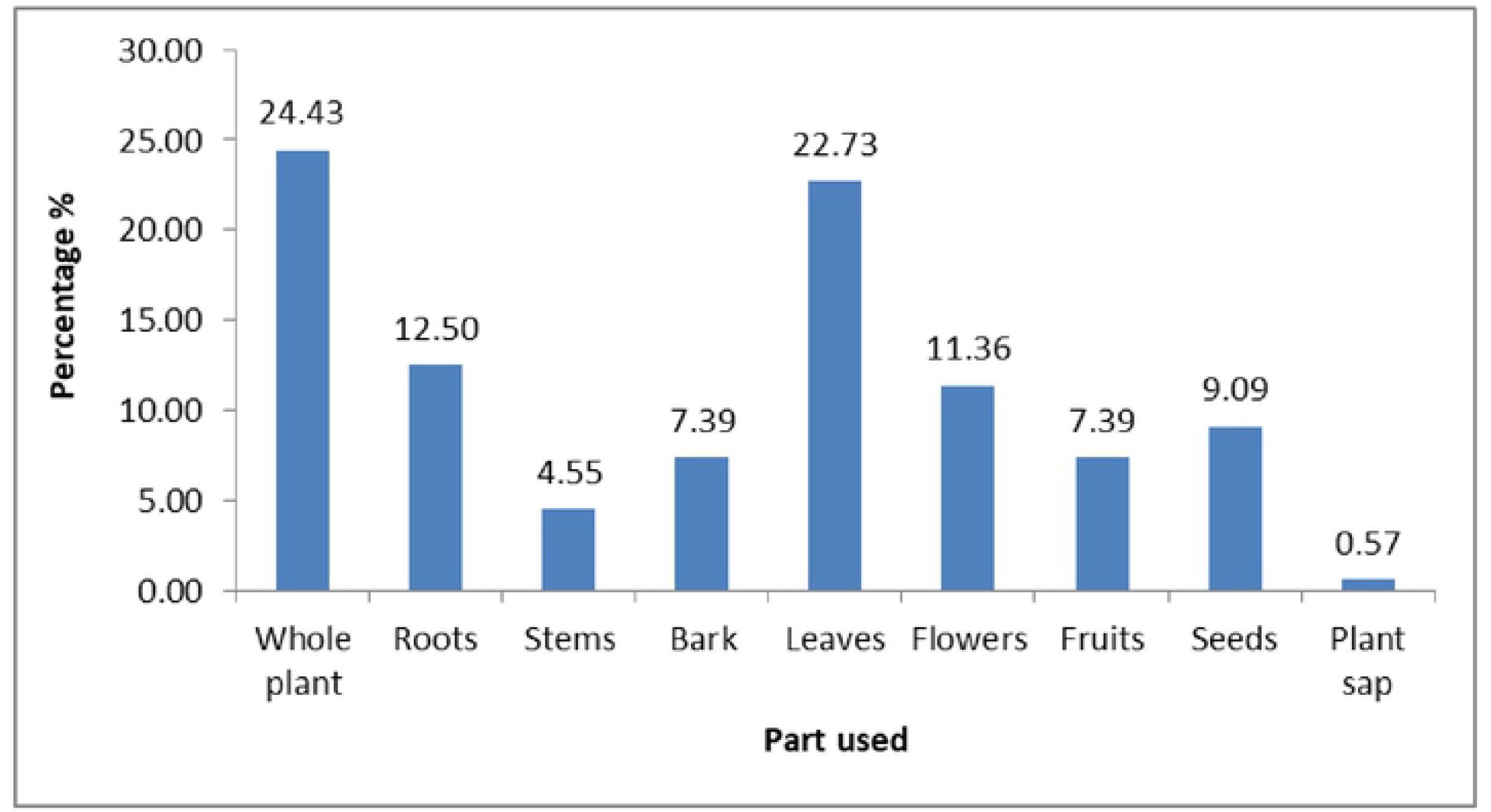
Most preferably used part for ethnomedicines

### Determination of ethnomedicinal value of enlisted shrubs on the bases of RFC (relative frequency of citation) and UV (used value)

In this study, the RFC ranged from 0.02 to 0.85. The highest RFC was found for *Rosa indica*, while lowest value was reported by *Deutzia scabra* and *Euonymus japonicus* (Table 3). However present study reported the UV from 0.01 to 0.23. The highest UV was found for *Jasminum grandiflorum* and *Hamelia patens*, while lowest was found for *Cestrum diurnum* and *Garcinia aristata.* Higher value of UV depends upon the number of used reports of plant species. Moreover all the listed ethnomedicinal shrubs were clustered in two ranking groups (i.e., high and low valued) on the basis of their calculated UV. Hierarchal cluster analysis on UV categorized the total 115 medicinal shrubs in two classes’ i.e. high valued ethnomedicinal shrubs and low valued ethnomedicinal shrubs (Table 4). The UV of high valued group ranged from 0.11 to 0.23 and this group comprised of 68 shrubs. However, the UV of low valued group of ethnomedicinal shrubs ranged from 0.01–0.09 with 47 species. Species under the label of high valued ethnomedicinal shrubs were recommended for further studies and can be used in development of drugs in the future. However, those species which reported as low valued ethnomedicinal shrubs were probably less famous or native people have less knowledge about them.

### Relative Importance (RI) and Fidelity Level (FL)

Number of diseases and number of body systems treated by a plant species is determined by RI value. In this study, it ranged from 0.16 to 43.73 (Table 3). The highest RI was accounted by *Tabernaemontana divaricata* while the lowest was found for *Garcinia aristata*. Fidelity level is use to determine the potential of one species to treat a particular disease. In this study, the FL ranged from 0.97% to 100. The plant species having the highest (100%) FL for treating specific disease was *Lawsonia inermis*, *Piper nigrum*, *Punica granatum*, *Rosa indica* and *Vitis vinifera* (Table 3). Plant have higher value of FL should be used in further herbal medicines.

### Informant Consensus Factor (ICF)

The ICF indicates the consensus between plants and informants in the treatment of diseases. Total diseases were categorized into 21 groups. ICF ranged from 0 to 0.45. The gastrointestinal diseases had highest ICF with 189 reports and 104 plants species. The second highest ICF (0.41) was found for respiratory diseases. Other studies have also reported the highest ICF for gastrointestinal tract diseases. The lowest ICF was found against blood purifier which showed that people were unaware about the use of shrubs related to certain diseases.

**Fig. 4.**
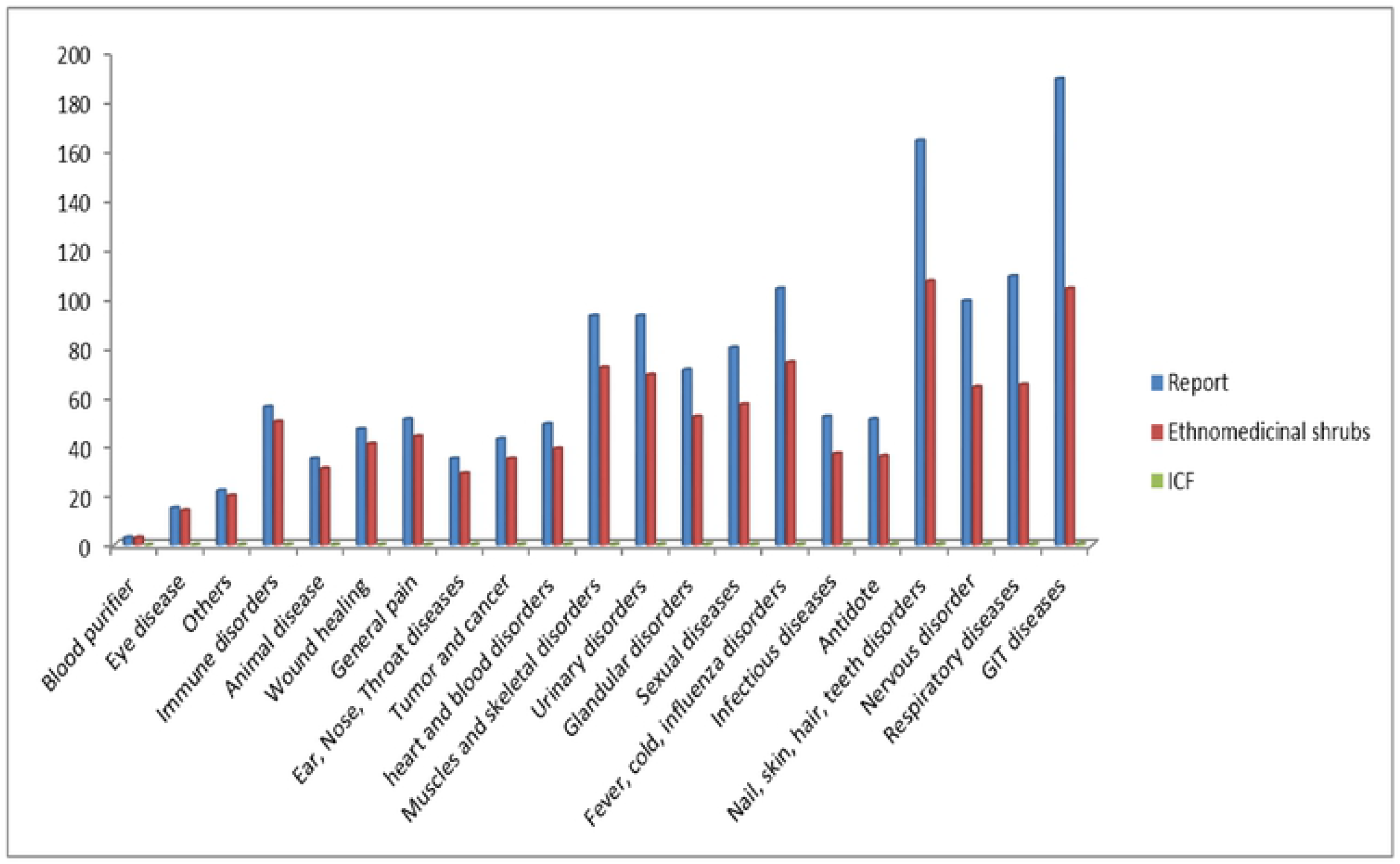
Summary of ICF

### Relative abundance of ethnomedicinal shrubs

In current study most of the ethnomedicinal shrubs were found to be rare in this study area i.e., 39.13%. However the relative abundance status of abundant, common and frequent ethnomedicinal shrub species were at the same level (12.17%). Moreover interesting relationship was observed by comparing the relative abundance categories and ranking groups of ethnomedicinal shrubs. It was found that maximum number of the high valued ethnomedicinal shrubs (n=29) were laid in the rare category of relative abundance which showed that priority of utilization of these ethnomedicinal shrubs directly affects their relative abundance. The shrubs of high valued group were more preferably used in comparison to the others, therefore their number declined due to over usage by the local people. Over utilization of medicinal species can lead to serious concern to plant conservation status and this fact was also supported by ethnomedicinal survey of FATA which revealed that some endangered species e.g., *C. tuberculata* and *N. ritchieana* due to over collection and utilization. Previous researchers also reported the loss of valuable medicinal plant due to various factors such as overgrazing, environmental degradation and deforestation. So there is a great need to conserve highly medicinal shrub species. Proper management and conservation of shrub species is needed.

**Fig. 5.**
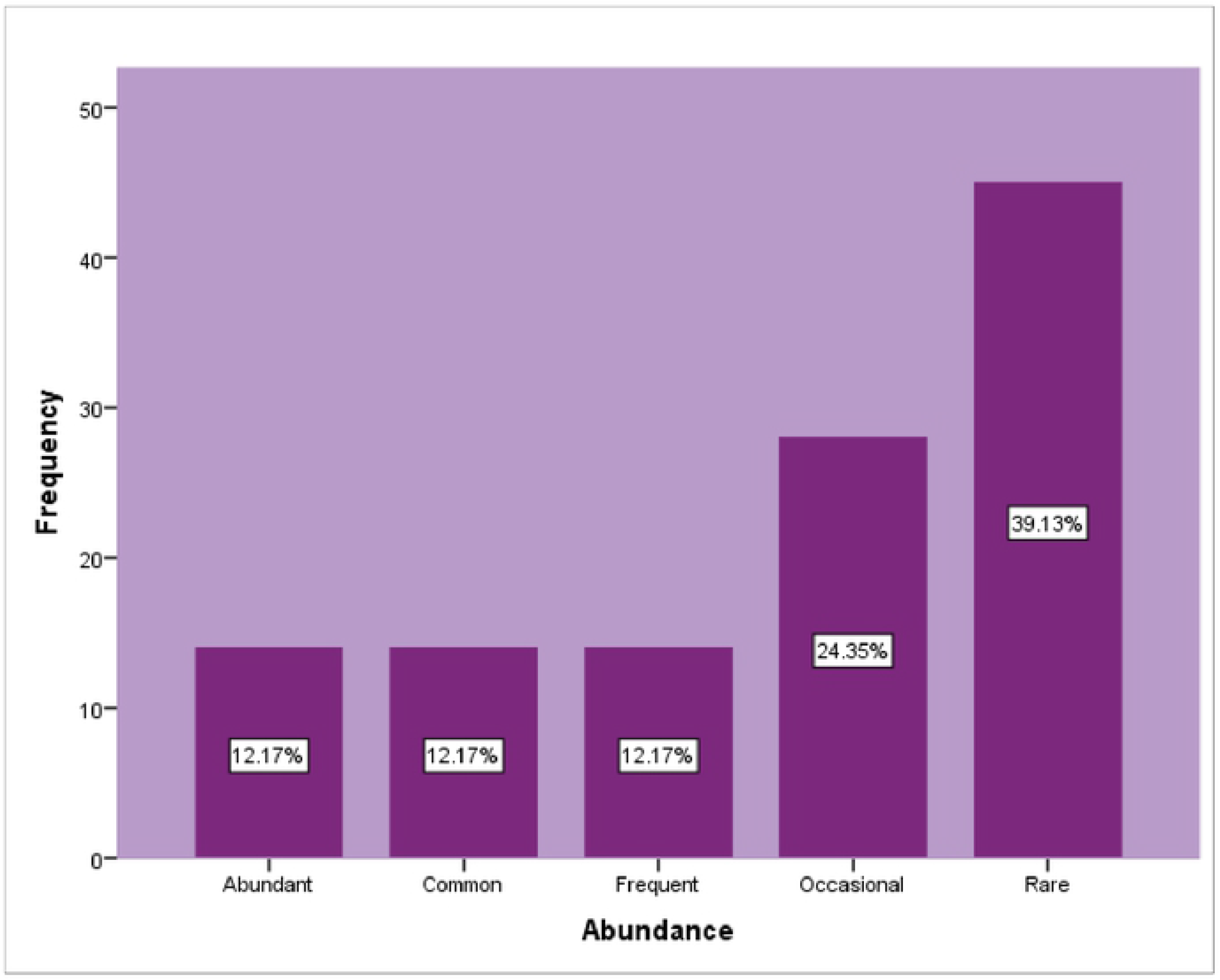
Relative abundance of enlisted ethnomedicinal shrubs

**Fig.6.**
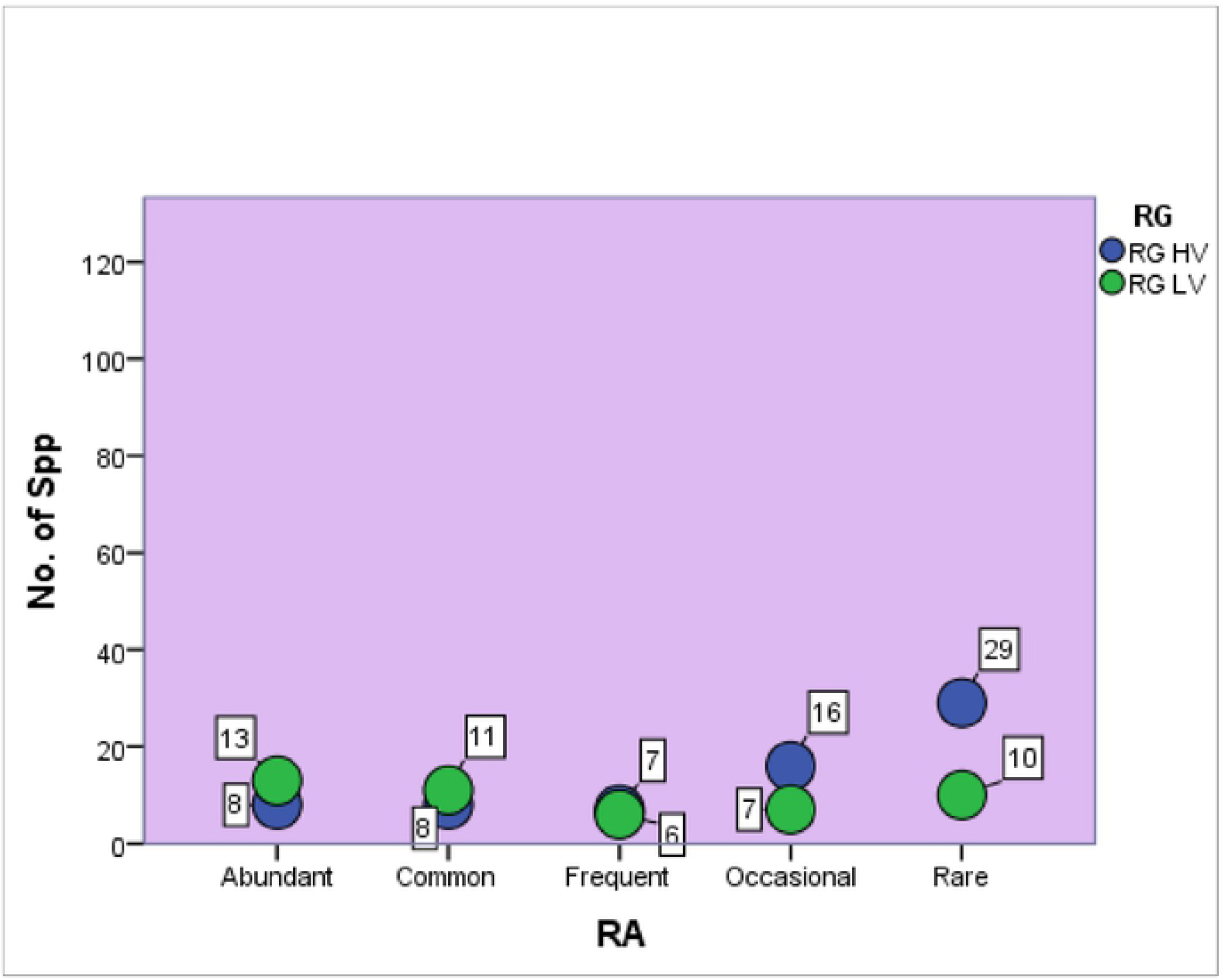
Association between abundance status and ranking groups of ethnomedicinal shrubs through cross tab analysis (SPSS 23)

### Ethnobotanical uses of listed ethnomedicinal shrubs

In the present study, ethnobotanical uses of one hundred and fifteen shrubs species were also documented which proved that shrubs provide multiple benefits such as fuel, wood for furniture, leaves and shoots used by animals and also provide a variety of products and services such as drugs, windbreaks, nectar sources, rubber, resin, gums, fiber, fencing materials, handicrafts, decorative material and oils. Such as *Lantana camara* is used for malarial treatment and related disorder*. Calotropis procera* is found to be effective for the treatment of gastrointestinal disorders. *Acacia farnesiana* has gummy roots which are used to cure sore throat and their flowers are involved in making ointment which is used to cure headaches. *Bauhinia tomentosa* is used to cure convulsions, constipation, sore throat and cough and its flowers are used for the remedy of diarrhoea and dysentery. *Salvia officinales* is used against cancer, in making drugs and is involved to deal with heart diseases. *Sambacus nigra* is used as laxative and to cure stomach disorders.

### Novelty and future impact

Current study is the first document on ethnomedicinal and ethnobotanical uses of 115 shrubs used by the inhabitants of district Lahore, (Punjab) Pakistan. In order to find the novelty index of current research the presently reported ethnomedicinal uses of shrubs were compared with previous ethnomedicinal studies conducted in similar study area(Table 1). Approximately, 9.5% similar ethnomedicinal shrubs were reported with more or less parallel uses; however, 90% shrubs were found new and had never been reported for their medicinal value (Figure 7). Out of 68 high valued ethnomedicinal shrubs 57 were reported first time for their medicinal uses in this study area. Some of these recently documented ethnomedicinal shrubs and their uses include: *Epipremnum aureum* (respiratory disorders), *Senna corymbosa* (asthma), *Sesbania sesban* (anthelmintic), *Tabernaemontana divaricata* (hypertension) *Tinospora sinensis* (arthritis) *Vitis vinifera* (skin allergies), *Salvia divinorum* (heart diseases), *Punica granatum* (dysentry), *Plumbago indica* (sore throat), *Opuntia ficus-indica* (laxative). These species with new medicinal uses could be further subjected for screening of bioactive compounds and their pharmacological activities. This can lead to introduce and develop novel drugs.

**Fig.7.**
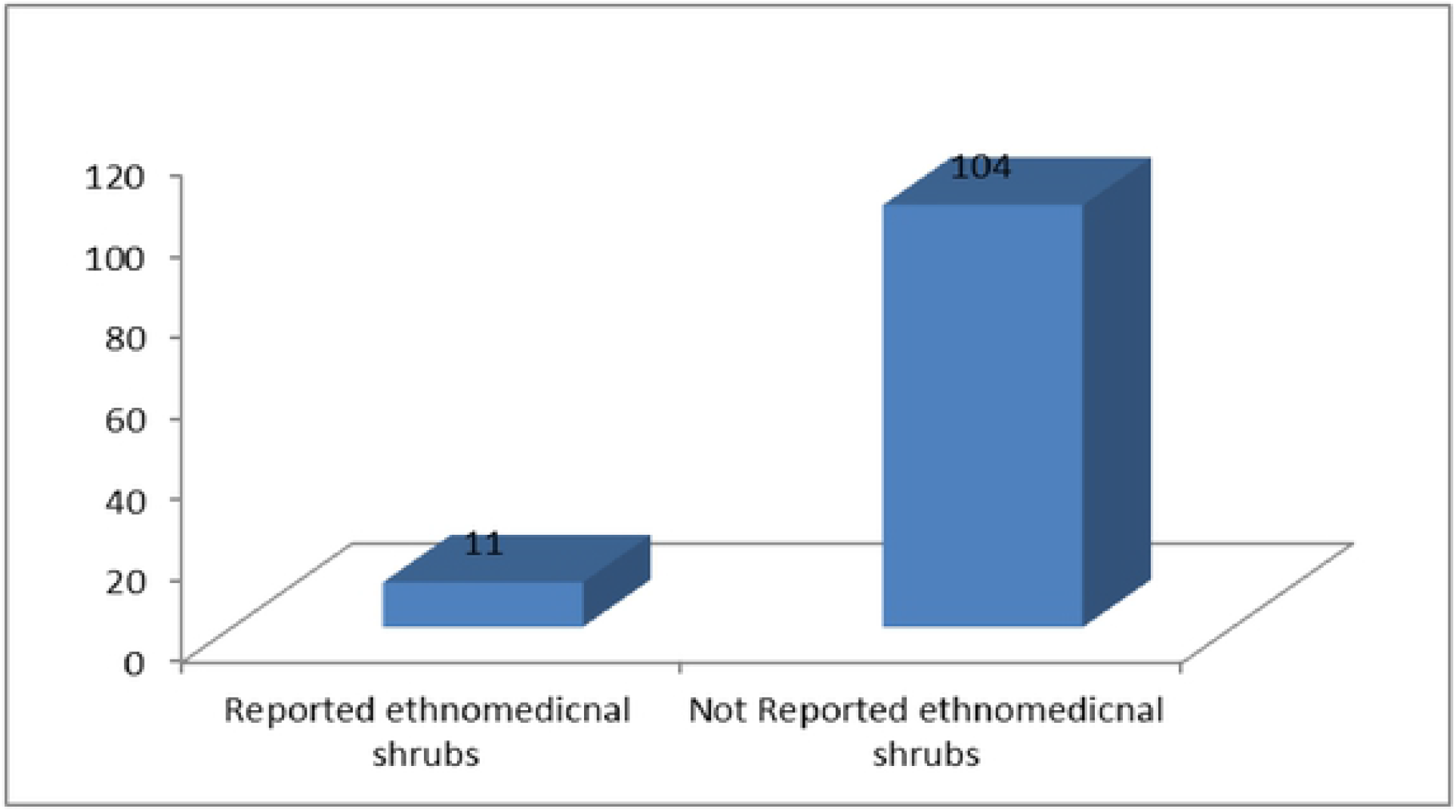
Novelty index

### Conclusion

Totally 115 ethnomedicinal shrubs were reported in this current research which reflects high diversity of medicinal plants in this study area possessed. This high medicinal plant diversity ultimately play valuable role in healthcare system of local people. It is essential to scientifically evaluate these ethnomedicinal shrubs for their pharmacological potential and toxicity profile. Furthermore the current endeavour not only contributed in preservation of traditional knowledge but also highlighted the conservation status of these medicinal shrubs. It was obvious form the present results that highly valued medicinal shrubs were decreasing day by day in number due to their over exploitation. Therefore serious precautions must be taken in order to preserve these natural resources. In other case ultimately it will affect the ethno-medical health care system and also depletion of raw materials will affect the discovery of potential drugs. Modern biotechnical methodologies such as genetic engineering, micropropagation and fermentation should be functional to enhance yield and modify the strength of medicinal species.

## List of abbreviations

RFC: Relative frequency citation
FC: Focal person count
N: total number of informants
SPSS: Statistical Package for the Social Sciences
RA: Relative abundance,
A: Abundant,
C: Common,
F: Frequent
O: Occasional
R: Rare
WP: Whole plant
Bk: Bark
FR: Fruit
LV: Leaves
FL: Flower
ST: Stem
RT: Root
PRE: Previously reported ethnomedicinal uses

## Declarations

### Ethics approval and consent to participate

Ethical approval was taken from the chairpersons of local government of all study areas of Lahore, Pakistan. Also we obtained permission from each informant before conducting the interview. However, all the participants were anonymised and so their personal details are not disclosed in this paper.

### Consent for publication

Not applicable

### Availability of data and material

Voucher specimens were submitted to the Herbarium of LCWU Pakistan for forthcoming uses (Table 3).

### Competing interests

Authors undoubtedly declared that they have no competing interests.

### Funding

No external funding resources were available for this particular study. All project costs were managed by either personally or resources of LCWU Pakistan.

### Authors’ contribution

The ethnobotanical survey and fodder grass sample collection was done by Shabnum Shaheen sobia sarwar, Iqra saeed, Nidaa Harun and Sana Khalid. The collected samples were well preserved and submitted in herbarium by Sehrish Sadia, Riffat Saddique and Hanan Mukhtar. Khadim Hussain and Muhammad Asaf Khan did the statistical analysis. Moreover Shabnum Shaheen Iqra saeed, sobia sarwar and Nidaa Harun wrote the manuscript by providing critical interpretation of the outputs. Shabnum Shaheen also supervised the whole study as well as helped along with Farah Khan in the identification of specimens.

## Acknowledgement

We acknowledge Dr. Mushtaq Ahmed and Dr. Muhammad Zafar, Department of Plant Sciences, Quaid i Azam University Islamabad for authorizing us to use their herbarium.

